# Reduced selection during sweeps lead to adaptive momentum on rugged landscapes

**DOI:** 10.1101/2024.04.08.588357

**Authors:** Clifford Bohm, Vincent R. Ragusa, Charles Ofria, Richard E. Lenski, Christoph Adami

## Abstract

Evolutionary theory seeks to explain the remarkable diversity and adaptability of life on Earth. Current theory offers substantial explanatory power, but it overlooks important transient dynamics that are prominent only when populations are outside equilibrium, such as during selective sweeps. We identify a dynamic that we call “adaptive momentum” whereby lineages with a selective advantage can temporarily sustain more deleterious mutations. This reduction in the strength of purifying selection allows populations to explore fitness valleys that are usually too costly to enter, potentially leading to the discovery of otherwise inaccessible fitness peaks. Using mathematical and agent-based simulations, we demonstrate adaptive momentum and show how periods of disequilibrium become windows of enhanced adaptation. Genetic exploration can occur during these windows without requiring mechanisms such as changing environments or complex landscapes. Adaptive momentum provides a simple potential explanation for bursts of rapid evolution observed in nature, including in pathogens such as SARS-CoV-2 and cancers. (152 words)

## Introduction

Traditional evolutionary models propose a predictable, measured rate of evolutionary progress known as gradualism^1^, characterized by a consistent distribution of time intervals between beneficial adaptations. In contrast, empirical observations often reveal evolutionary trajectories that deviate from this predictability, presenting patterns that are less consistent and more varied than traditional models suggest^2,3^. Some theories associate these deviations in evolutionary rates with environmental change^4^ or complex interactions between mutations^5^. However, in cases where these effects are not evident, such theories may not satisfactorily explain observed evolutionary patterns. Here, we propose an alternative model that predicts varied rates of evolution while relying on purely Darwinian mechanisms. Our model complements, rather than replaces, these existing theories by offering a different perspective for understanding evolutionary dynamics in specific contexts.

We argue that traditional population-genetic models have over-emphasized the equilibrium dynamics of stable populations, thereby minimizing the importance of transitional periods of disequilibrium that inevitably arise during selective sweeps, range expansions, and other similar phenomena. Although these transitional periods typically constitute only a small fraction of evolutionary timelines, our findings illustrate their potential to leave lasting imprints on evolutionary trajectories.

Incorporating insights from foundational evolutionary theories, we highlight dynamics that, although often dismissed, become pronounced during these transitional disequilibrium periods. Specifically, we introduce the theory of “adaptive momentum,” which we define as a temporary reduction in selection against deleterious mutations that results from fitness disequilibrium during these periods. This reduced selection can allow increased mutational buffering, because some mutations are shifted from being deleterious to “nearly neutral”^6^, resulting in the persistence and accumulation of these mutations via hitchhiking. Consequently, there is more genetic exploration^7–10^, and thus increased potential for the discovery of beneficial mutations across fitness valleys^11,12^. Critically, adaptive momentum explains why populations that have recently experienced a disequilibrium-inducing event are likely to experience more change while the disequilibrium persists.

The disequilibrium states that allow for adaptive momentum can be initiated by the spread of a beneficial mutation, environmental change, range expansion, or increased carrying capacity. Regardless of the event that causes disequilibrium, the ensuing period should promote exploration before stabilizing at a new equilibrium. By focusing on transitional dynamics, adaptive momentum provides a novel framework that bridges microevolutionary processes and macroevolutionary patterns, thereby enabling a more comprehensive understanding of the evolutionary process.

## Conceptualization

We illustrate the theory of adaptive momentum via selective sweeps in Fig. 1. The blue “water line” depicts the mean fitness of an asexual population. Any phenotypes with fitness above this line will tend to increase in frequency, even if they do not have the highest fitness relative to every other phenotype^13^. In Fig. 1a, a population centered on fitness peak *x* has just discovered a new higher peak *y*. During the ensuing selective sweep, individuals centered on the higher peak experience adaptive momentum because they have relatively higher fitness versus the average population fitness, even if some of them have acquired deleterious mutations that reduce their fitness below the new, higher peak. This higher-than-average fitness facilitates increased genetic exploration, allowing them to enter fitness valleys and potentially cross them, as seen in Fig.1b. If the population does not traverse another fitness valley before fixing on the new peak, the scenario depicted in Fig. 1c unfolds. Here, the valley becomes “submerged” as the average fitness increases, and mutations that had been buffered during the disequilibrium phase are purged, making it, once again, highly unlikely for the population to cross the fitness valley. Conversely, if a path to greater fitness is discovered before the population becomes fixed on peak *y* (Fig. 1d) then a new selective sweep begins, extending the period of adaptive momentum.

**Fig. 1.**
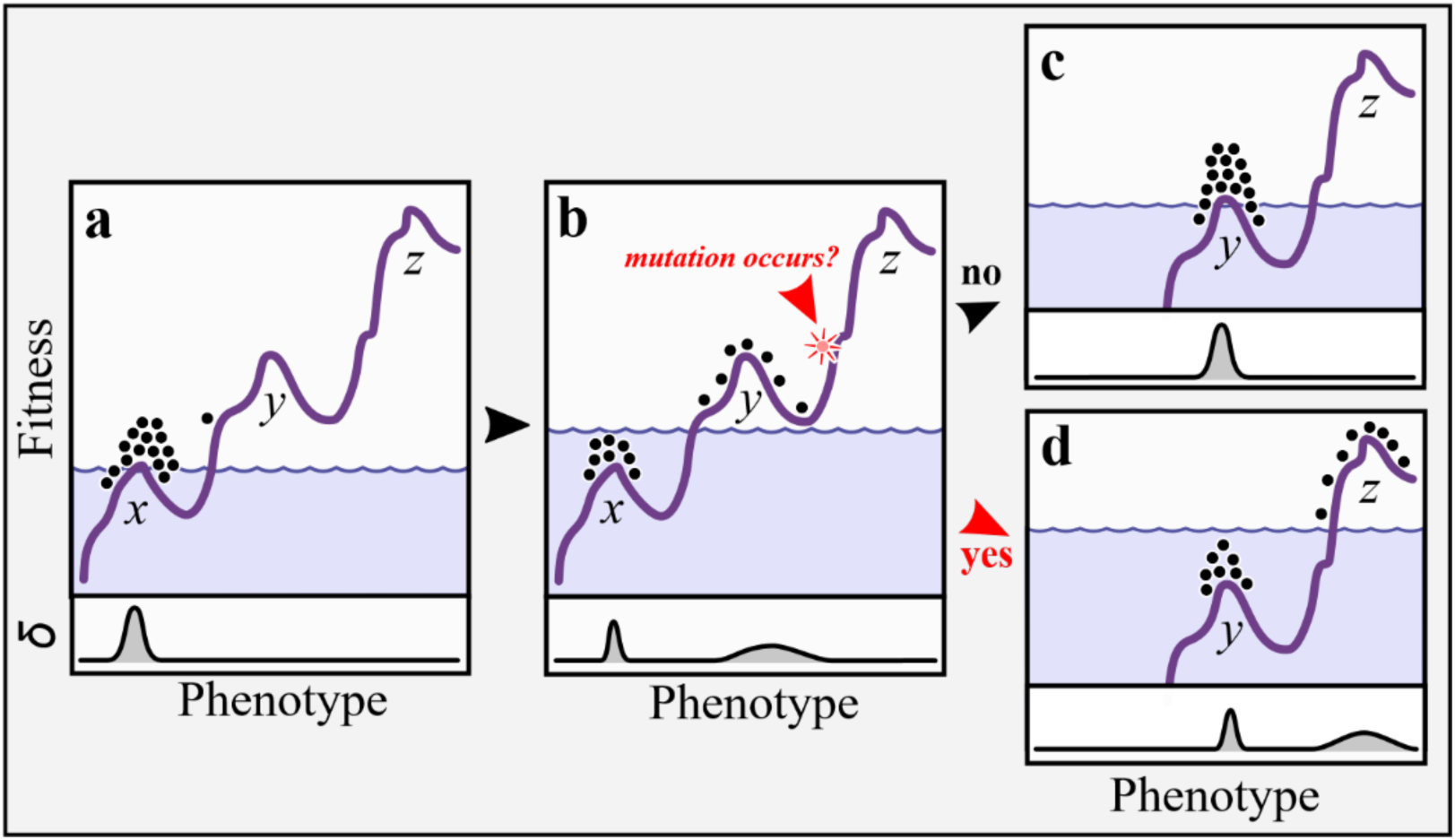
Adaptive momentum in an asexual population evolving on a multi-peaked fitness landscape. Arrows between panels indicate temporal progression, dots indicate extant phenotypes in the population, and the blue “water line” shows the population’s average fitness. The bottom of each panel shows a phenotype density plot. **a**, A sweep has just started in the population. Most individuals are centered on peak *x* with a single individual on the slope leading to peak *y*. **b**, The selective sweep has progressed. Those individuals near peak *y* have above-average fitness and thus experience reduced purifying selection. The resulting increase in diversity increases the probability that a mutant will cross the fitness valley to the base of peak *z* (red arrow). **c**, If such a mutation does not occur, the average fitness will increase as purifying selection tightens the distribution of phenotypes around peak *y*. The adaptive momentum will be lost, however, reducing the probability of finding peak *z*. **d**, If such a mutation does occur, a second peak shift (this time to *z*) is initiated. The population may subsequently fix on the new peak, or it may continue its momentum by making additional genetic discoveries.

We define a “momentum window” as the period during which adaptive momentum dampens the strength of selection against deleterious mutations. This window opens with the onset of disequilibrium and closes when an equilibrium is reestablished. While the window is open, the reduced purifying selection facilitates crossing fitness valleys. Clusters of valley crossings during a single momentum window are termed “cascades”.

When such cascades are interspersed with prolonged periods of stasis, it leads to a pattern of punctuated equilibrium^2^ and generates dynamics that correlate with the over-dispersed molecular clock model^14–16^, as discussed below.

If a fitness landscape is mostly neutral or contains many beneficial mutations, the increased exploration from adaptive momentum may not improve adaptation, since adaptation does not require crossing valleys. Moreover, in some landscapes, adaptive momentum could even impede adaptation to landscapes rich with beneficial mutations. On such a landscape, the reduction in selection afforded by adaptive momentum not only increases the potential for exploration, but also decreases the effectiveness of selection at promoting beneficial mutations within an advantaged clade. This suggests that in landscapes with many beneficial paths, momentum windows can lead to periods of inefficient genetic search. There is a tradeoff: while adaptive momentum during transitions can enhance exploration, it does so at the cost of effective exploitation.

The evolutionary regime^17,18^ plays a pivotal role in determining the effects of adaptive momentum on a population. At one end of the spectrum, the *sequential fixation* (SF) regime describes a condition where each beneficial mutation tends to fix before another can be discovered. In this regime, any momentum windows are generally too short to allow the accumulation of the deleterious mutations needed for valley crossings. At the other end, the *clonal interference* (CI) regime describes the condition in which new beneficial mutations arise more rapidly than they can fix. In this regime, the rate of adaptation is limited by the rate of fixation rather than the beneficial mutation rate; the rate of adaptation cannot be predicted simply by the discovery rate because most beneficial mutations are lost by clonal interference. Although adaptive momentum intensifies exploration in the CI regime, the increased rate of discovery is overshadowed by the loss of those beneficial mutations that fail to fix^19,20^.

Both the SF (strong selection/fitness) and CI (clonal interference) regimes tend to mitigate the impacts of adaptive momentum on evolutionary outcomes. However, there exists a ‘balanced regime’ between them, characterized by roughly comparable fixation and discovery times. In this regime, adaptive momentum can significantly enhance adaptation rates, as it allows for additional discoveries during momentum windows without being overburdened by clonal interference.

To demonstrate the influence of adaptive momentum on evolutionary dynamics, we conduct two main experiments. In the “three-peaks” experiments, we examine a system in the balanced regime to demonstrate the difference between populations experiencing adaptive momentum and populations at equilibrium. In the “infinite peaks” experiment, we explore the evolutionary dynamics of populations on an unbounded fitness function to display behaviors associated with SF, CI, and the balanced regime. We use these results to explain how the mode of evolution impacts adaptive momentum, leading to altered rates of evolutionary discovery.

## Results

### Evolving populations have greater potential during disequilibrium

Given a population at equilibrium, we can calculate the expected time for that population to discover a path to higher fitness through a given valley. However, during a period of adaptive momentum, the population is not in equilibrium, and the mathematical models that describe equilibrium populations do not apply.

Using the “three-peaks” simulation, we demonstrate that populations experiencing adaptive momentum cross fitness valleys more readily than populations at equilibrium, even if the population at equilibrium initially has a higher average fitness. This result shows that the dynamics of valley crossing can be altered by adaptive momentum. The simulations use a simple three-peak fitness function (see Fig. 2a), and we tested two different initial conditions for the populations.

**Fig. 2.**
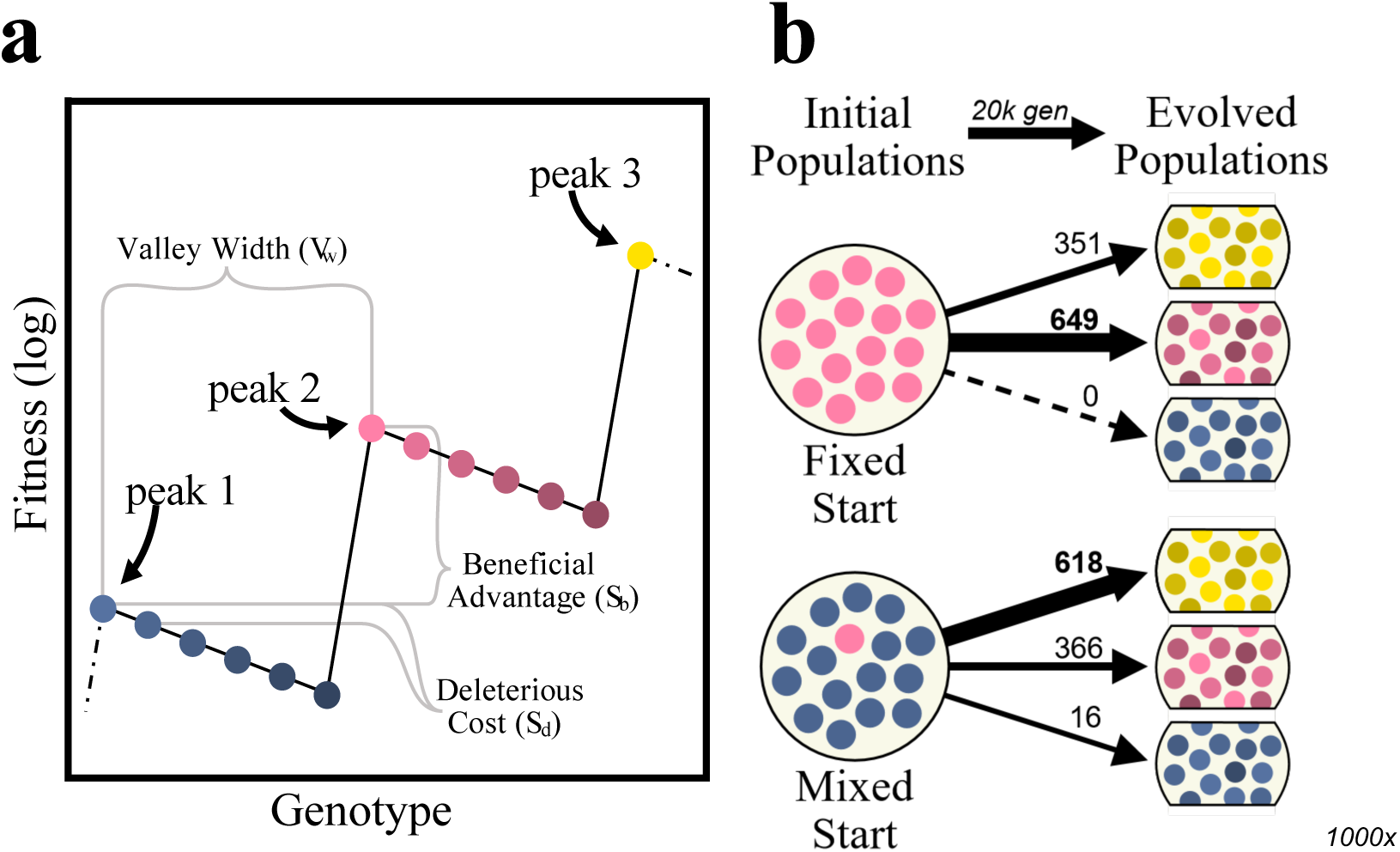
The “three-peaks” experiment. **a**, Fitness function for the “three-peaks” experiment, color-coded based on fitness. **b**, Number of replicate experiments (out of 1000) that reached the respective peaks after 20,000 generations, given starting conditions where populations are either initialized on peak 2 (Fixed Start) or have only just discovered peak 2 (Mixed Start) and are thus experiencing adaptive momentum.

In the “fixed start” condition, a homogeneous population of 2400 individuals started on the middle peak (peak 2 in Fig. 2a), simulating a population initially at equilibrium. In the “mixed start” condition, we initialized a population with the majority of individuals (2395) on the lowest peak (peak 1) and five individuals on peak 2, simulating a population that has just begun a genetic sweep (and therefore is out of equilibrium). We tested each initial condition 1000 times and recorded the highest peak reached at the end of each test (see Methods for details.)

The results in Fig. 2b show that only 35.1% of the populations in the fixed-start condition crossed the fitness valley to reach peak 3, whereas in the mixed-start condition 61.8% reached peak 3. This difference occurs despite the initially higher average fitness of populations in the fixed-start condition (all individuals on the higher peak 2) than in the mixed-start condition (only ∼2% of the population on peak 2). Extended Data Fig. 1 show results of the same experiment using other population sizes and different values for the valley depth. These results show that the effects of adaptive momentum are lessened if either rapid fixation or clonal interference dominate the dynamics, as we discuss in more detail below.

### Adaptive momentum depends on the relation between discovery and fixation times

The “three-peaks” experiment demonstrates a clear difference in the adaptive potential of populations in equilibrium and disequilibrium states. However, it does not explain the underlying cause of these differences. Specifically, it does not identify the evolutionary parameters and mechanisms that account for the differences. To provide a more detailed exploration of adaptive momentum and its potential to influence evolutionary rates, we extended the fitness function in the “three-peaks” experiment to create the “infinite-peaks” experiment.

We identify four quantities that are crucial for understanding adaptive momentum. The *observed crossing time* (*T*_obs_) is the average time required for an evolving population to cross fitness valleys and establish at a new peak when the population is allowed to run for many uninterrupted generations. The *stochastic tunneling time T*_st_ ^21^ is the expected time for an equilibrium population (i.e., starting on an adjacent but lower peak) to discover a series of mutations that allow it to cross a single fitness valley and establish at a higher peak across the valley. The *neutral drift time T*_nd_ represents the time it takes to cross the same valley and establish if the intervening mutations were neutral instead of deleterious; *T*_nd_ is a lower limit on *T*_st_. Finally, the *fixation time T*_fix_^22,23^ quantifies how long, on average, it takes a valley-crossing beneficial mutation to fix in a population after its origination (assuming the mutation does fix, as opposed to being lost due to drift). See Extended Fig. 2 for schematic representations of these quantities.

To examine the effect of adaptive momentum across different modes of evolution, we conduct experiments under conditions ranging from the SF regime (where *T*_fix_ ≪ *T*_st_) to the CI regime (where *T*_fix_ ≫ *T*_st_). We use population size to vary the relation between *T*_st_ and *T*_fix_, although other parameters, such as mutation rate or valley depth and width, can be used as well (see Extended Data Fig. 3 and Extended Data Fig. 4 for additional results.)

Fig. 3a shows the mean values of *T*_obs_, *T*_st_, *T*_nd_, and *T*_fix_ as a function of population size. Note that population sizes are shown on a logarithmic scale. At small population sizes, the results are consistent with the sequential fixation regime, namely *T*_obs_ tracks *T*_st_. In this regime, *T*_fix_ ≪ *T*_nd_ and consequently, individual beneficial mutations fix more quickly than new mutations are discovered. At the large population extreme where *T*_fix_ ≫ *T*_st_, the dynamics are dominated by clonal interference (the CI regime). There, *T*_obs_ *> T*_st_ because new beneficial mutations are often discovered before previous ones can fix. In this regime where discovery is relatively quick, beneficial mutations are likely to arise in different clades and compete for dominance. In this case, the factor limiting the rate of adaptation is not the rate of discovery, but rather the rate of fixation.

**Fig. 3.**
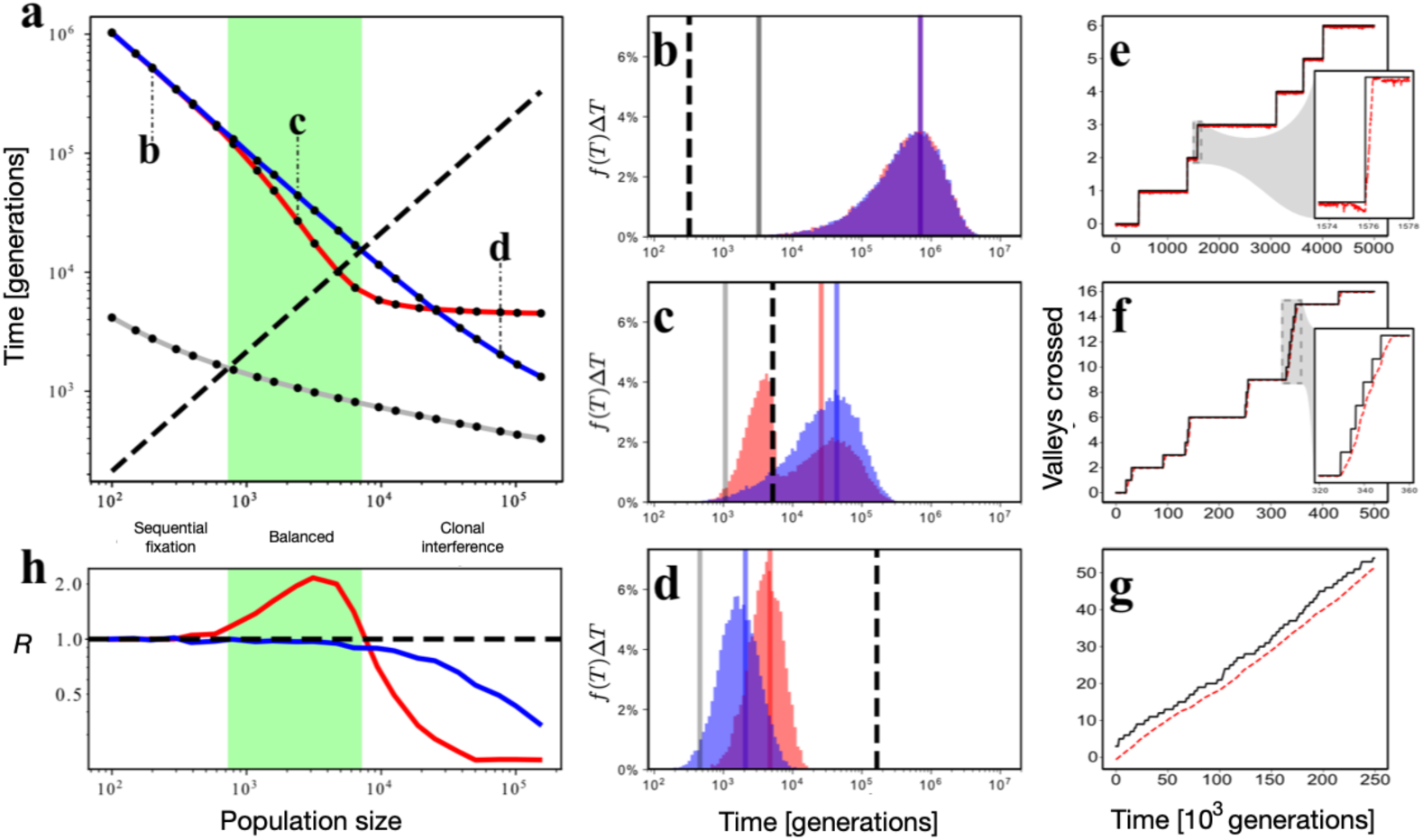
Effects of population sizes on valley crossings and fixation time in the “infinite-peaks” experiment. **a**, Stochastic tunneling time (*T*st) in blue, neutral drift time (*T*nd) in gray, fixation time (*T*fix) in black, and the observed crossing time (*T*obs) in red as a function of population size for the fitness function described in Fig. 2a, except with an infinite series of valleys and peaks. Dots are averages of collected data (see Methods), with 95% confidence intervals smaller than the diameter of the dots. The labels **b**, **c**, and **d** in panel **a** refer to population sizes characteristic of the SF, balanced, and CI regimes respectively. Panels **b** through **d** show the distributions *f*(*T*) for *T*st and *T*obs (see Methods) whose averages are shown in **a** at the indicated population sizes, with vertical lines showing those averages along with the averages for *T*nd and *T*fix for reference. Panels **e** through **g** show typical evolutionary trajectories over arbitrarily chosen periods for the population sizes in **b** through **d,** respectively. The black line is the highest log fitness in the current population (i.e., maximum peak discovered) and the red-dashed line is the average log fitness. The difference between the black and red-dashed lines is only clearly visible in **g**; inset plots have been added to **e** and **f**. In all experiments, populations evolved on a periodic fitness landscape with valleys defined in Methods, Eq. (3), using *Vw* = 6, *Sb* = 0.5, *Sd* = 0.03, and mutation rate *µ* = 0.0025. Panel **h** shows the index of dispersion *R* calculated from the observed valley-crossing times *T*obs (red line) and the stochastic tunneling times *T*st (blue line). The dotted line at *R* = 1 is the expectation for a Poisson process, with the mean equal to the variance. Population sizes associated with the three evolutionary regimes are labeled, illustrating that the balanced regime (green shaded area in **a** and **h**) coincides closely with population sizes where the observed transition times are reduced, indicating accelerated valley crossings.

Between the small and large population sizes, where *T*_fix_ is between *T*_nd_ and *T*_st_, we observe a “balanced” regime. Here, the observed valley-crossing time *T*_obs_ is substantially shorter than the expected crossing time by stochastic tunneling, *T*_st_. This difference occurs because the discovery time in the balanced regime is short enough relative to the fixation time to reduce the impact of clonal interference, but long enough to allow sufficient accumulation of deleterious mutations (during a momentum window) to enable populations to cross adaptive valleys.

Figs. 3b-d show the distributions *f*(*T*) behind the averages shown in Fig. 3a for three population sizes that correspond to the distinct adaptive regimes. Fig. 3b shows the distributions at *N* = 200, Fig. 3c at *N* = 2,400, and Fig. 3d at *N* = 76,800. These data show key elements that give rise to the dynamics of adaptive momentum. The distributions of *T*_st_ are unimodal in all three cases, with averages that decrease as the population size increases. Conversely, the *T*_obs_ distributions are neither always unimodal (Fig. 3c), nor do they exhibit as consistent a relation with population size as observed with *T*_st_. At *N* = 200 (Fig. 3b), the close match between the distributions of *T*_st_ and *T*_obs_ supports the assertion that observed valley crossings occur by stochastic tunneling in the SF regime. By contrast, at *N* = 76,800 (Fig. 3d) the *T*_obs_ and *T*_st_ distributions have different means. The positive offset between *T*_obs_ and *T*_st_ shows that, although discoveries of new peaks can occur rapidly in this regime, some of the discoveries are lost because of clonal interference (see Fig. 4 for an illustration of this effect).

**Fig. 4.**
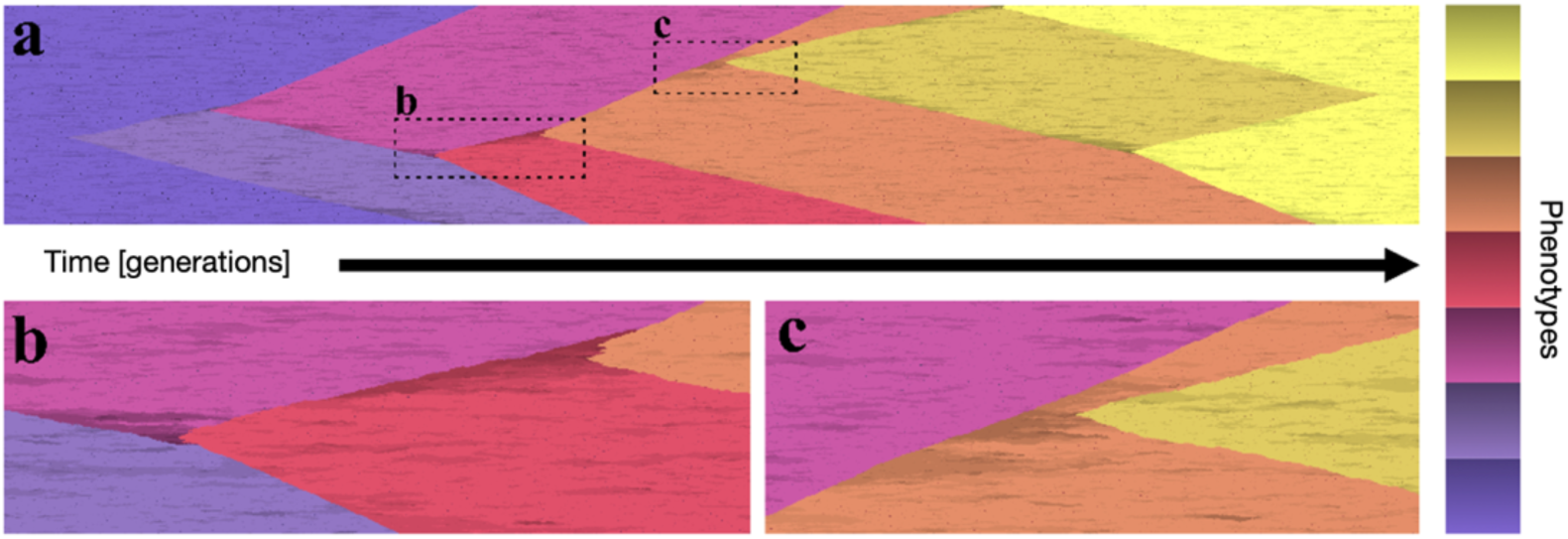
Visualization of adaptive momentum. **a**, A population of 4,800 individuals on a one-dimensional strip evolved for 31,000 generations, with the fitness function and other parameters as in Fig. 3. Each saturated color corresponds to one peak (color scale on right), with darker shades indicating lower fitness genotypes in the adjoining valley. Generations advance from left to right. Valley crossings (darker hues in the background of the preceding type) lead to the discovery of new peaks, which give rise to expanding segments with the new colors. **b**, Detail of the second and third crossing events showing how the beneficial mutations that discover new peaks occur at the edge between the sweeping and background clades, where deleterious mutations that cause fitness deficits (dark hues) are partially shielded from purifying selection. **c**, Detail of the fourth crossing event in the cascade, where a beneficial mutant is generated further from the boundary between the clades, but again along a line of individuals with impaired fitness (darker hues).

Between these extremes, at *N* = 2,400 (Fig. 3c), the distribution of observed discovery times is bimodal: one mode of the *T*_obs_ distribution aligns with the distribution of *T*_st_, while the other mode consists of discovery times that that are *shorter* than *T*_fix_. These data indicate that *two different processes* are giving rise to the valley crossings, with different associated rates of adaptation. The mode corresponding to *T*_st_ is composed of discovery times from periods of equilibrium and associated stochastic tunneling across the valleys, while the mode with shorter discovery times indicates adaptive momentum with clusters of valley crossings during momentum windows. Moreover, the mode consisting of shorter observed times is bounded by the fixation time (*T*fix, vertical dashed black line), which reinforces the fact that the discovery of new peaks in this mode must occur before fixation (i.e., within a momentum window).

Figs. 3e-g show typical examples of evolutionary trajectories (log fitness over time) that illustrate valley crossing dynamics for the three different regimes in Figs. 3b-d. Each increase in the black line corresponds to the discovery of a new peak. The red-dashed line shows the average log fitness of the population, which defines the “water line” in Fig. 1. The population is in disequilibrium whenever the red-dashed line is substantially below the black line, which indicates that the population is shifting between peaks and experiencing adaptive momentum. For small populations in the SF regime (Fig. 3e), fixation occurs rapidly relative to the time needed to discover a new peak (see inset), leaving only a short momentum window. For large populations in the CI regime (Fig. 3g), average fitness consistently lags behind the highest current fitness because discoveries of new peaks occur before an individual valley-crossing mutation can fix. Thus, while adaptive momentum is essentially constant in sufficiently large populations (i.e., the momentum window never closes), the observed rate of adaptation (*T*^$%^) is still smaller than what the stochastic tunneling time (*T*_st_) would predict, owing to clonal interference. Fig. 3f shows how, for an intermediate population size within the balanced regime, adaptive momentum results in cascades of valley crossings to higher peaks that alternate with periods of stasis, giving rise to punctuated evolution. The inset shows how a momentum window is kept open by each discovery of a new peak during such a cascade. The window eventually closes when the population fixes on a peak before another discovery is made, as evidenced by the average fitness (red-dashed line) catching up to the maximum fitness (black) and remaining there for some time. Fig. 3f is discussed in the section “Adaptive momentum produces an over-dispersed clock” below.

In both the three-peaks and infinite-peaks experiments, the populations evolved in a physically structured environment, with individuals arrayed along a one-dimensional ring. This approach is both computationally efficient and generates results that clearly illustrate the dynamics under investigation. In Extended Data Fig. 5, we show that adaptive momentum also occurs in well-mixed populations, although the effect is more subtle and requires larger population sizes. We discuss the effects of population structure and the well-mixed results shown in Extended Data Fig. 5 in the Supplementary Discussion, sections “Adaptive momentum is modulated by population structure” and “Adaptive momentum in unstructured populations” respectively.

### In spatially structured populations, valley crossings tend to occur on the leading edge of selective sweeps

The role of adaptive momentum in propelling populations across fitness valleys is most easily visualized in a one-dimensional population, where the fitness of every individual is shown over time and the locations of innovations and selective sweeps are visually prominent (see Methods). Fig. 4 shows a typical example of an evolutionary history in the balanced regime, illustrating a cascade with six sequential crossings. Here, time (in generations) progresses from left to right, where each column depicts the fitness of an individual in the population at that moment and location. Each hue represents a different peak, with darker shades of that hue depicting lower-fitness genotypes in the next valley. Because all transitions to higher peaks must occur through the valley that precedes it, we only see a sharp change in hue if the immediate progenitors of that transition exhibit a dark shade of the preceding hue. Each valley crossing (excluding the first) occurs at the leading edge of a selective sweep, where there are lineages in which successive generations experience reduced selection owing to adaptive momentum. Note that the last peak (yellow) was independently discovered twice (i.e., at two distant locations). Furthermore, the slope of each leading edge indicates how quickly each type is expanding. Thus, the growth rate increases (and the fixation time decreases) with the magnitude of the fitness differential at the interface between types. Figures 4b-c provide a closer look at the transitions that allow valley crossings, highlighting the accumulation of deleterious mutations at the leading edge of sweeps.

### Adaptive momentum produces an over-dispersed clock

Kimura’s neutral theory^24^ predicts that the accumulation of neutral mutations (i.e., those that are neither beneficial nor harmful) should follow a Poisson distribution. In this context, the dispersion coefficient *R* is a key metric, quantifying the variance of a distribution relative to its mean. A Poisson distribution of events results in unit *R*, consistent with a process producing independent events. Deviations of *R* from 1 are indicative of processes that are either non-random or non-independent. Under-dispersion (*R <* 1) suggests a more regular occurrence of events, while over-dispersion (*R >* 1) points to bursts of events, suggesting punctuated dynamics (as in “punctuated equilibrium”). In the context of neutral theory, the latter phenomenon has been termed an over-dispersed molecular clock^15^. It is important to note, however, that it is the *independence* of the events, and not the neutrality of mutations *per se*, that results in *R*=1. As a consequence, the relevant events for the purposes of this paper are not mutations; instead, the events are the valley crossings that lead to the discovery of new fitness peaks.

In our model, the accumulation of beneficial mutations in *T*_obs_ correspond with these dispersion patterns. Recall that Figures 3b-d show the observed distributions of times between events for populations under the SF, balanced, and CI regimes, respectively. However, the dispersion calculation requires counts of events per unit time. Therefore, we must first convert the “times-between-events” data into “events-per-unit-time” data (see Methods section), which then allows us to calculate the dispersion coefficient of these new distributions, presented in Figure 3h. For conditions in the SF regime, where *T*_st_ ≪ *T*_fix_, we observe that *R* is approximately one. Although the mutations are driven to fixation by selection in this regime, the resulting “adaptive clock” depends on the temporal independence of the valley crossings. The clonal interference regime where *T*_st_ ≫ *T*_fix_ results in an *R < 1*, consistent with existing literature^15,25–27^. Importantly, in the balanced regime, when *T*_nd_ ≤ *T*_fix_ ≤ *T*_st_, we see *R > 1.0*, indicating an over-dispersed adaptive clock. This finding suggests that, in addition to the above relations between *T*_st_, *T*_nd_, and *T*_fix_, a dispersion of adaptive events greater than one is another indicator of the balanced regime.

### Is adaptive momentum important in natural systems?

At a fundamental level, adaptive momentum is an unavoidable feature of the process of adaptation in populations, because the dynamical spread of beneficial mutations reduces the effective strength of selection against deleterious mutations in the lineages that have beneficial mutations. This weaker selection against deleterious variants thus allows populations to explore lower fitness regions of the adaptive landscape, and potentially to cross valleys to peaks with higher fitness. Thus, adaptive momentum is most important when evolution occurs on a rugged, multi-peaked fitness landscape, such as those used in our models. The form and ruggedness of real fitness landscapes is, unfortunately, poorly documented in natural systems^28^. However, it is widely accepted that these landscapes must be multi-peaked on some scale to account for the diversity of life. This realization has led to the much-debated question of *how* populations can cross adaptive valleys and thereby discover new peaks.

We expect that natural populations sometimes experience cascades of adaptation because of adaptive momentum. These cascades might be rare, but when they occur, they would have an outsized effect in determining the ‘winning’ lineages over time. However, it can be difficult to disambiguate whether a period of rapid adaptation is due to adaptive momentum or other effects, such as a response to environmental change or trait interactions. Specifically, it is challenging to find conclusive evidence that a given line of descent has crossed a fitness valley that could not have been crossed without adaptive momentum. Doing so would require both a detailed knowledge of all possible evolutionary pathways on the fitness landscape as well as confirmation that there were no external factors (such as environmental change) that could account for the changes in selective pressure (and concomitant rapid adaptation).

Although detecting the effects of adaptive momentum may be difficult in practice, it is easy to imagine important scenarios where adaptive momentum could play a major role. For example, in the origin and spread of many cancers, multiple somatic mutations appear in rapid succession. It is reasonable to hypothesize that in this non-equilibrium setting, one or more beneficial “driver” mutations^29^ generate adaptive momentum that enables the rapid accumulation of additional mutations, resulting in a more aggressive cancerous cell. Similarly, adaptive momentum may play a large role in the evolution of viral populations. In particular, a pandemic might start with a beneficial mutation that increases a pathogen’s transmission in a new host, resulting in a selective sweep that allows deleterious mutations to hitchhike with the successful lineage as it spreads. In the case of SARS-CoV-2, for example, multiple waves of variants have appeared and swept through the global population. Each wave creates a new opportunity for adaptive momentum, and some important new variants differ from their progenitors by tens of mutations. These facts indicate rapid accumulation and fixation of multiple mutations in certain lineages, which then rise to high frequency globally. The identification of clusters of unusual, rare, and deleterious mutations at previously highly conserved sites of the SARS-CoV-2 Spike protein^30^ in the BA.1 variant suggests multiple valley-crossings in close succession. And indeed, there is strong evidence for an over-dispersed clock during SARS-CoV-2 sweeps^31^.

On a geological timescale, the fossil record documents several periods of rapid change including the fossils of the lower and middle Cambrian^32^, phanerozoic marine fossils^33,34^, and tracheophyte (vascular land plant) fossils^35,36^. While the absence of genomic data makes it challenging to determine the precise role of adaptive momentum in these cases, several factors suggest that adaptive momentum could have played a significant part. Although recombination in sexually reproducing lineages can disrupt the epistatic combinations of mutations that promote adaptive momentum, physically linked blocks of genes in disequilibrium may still provide the necessary variation for this process. This is particularly likely in structured populations with loosely connected and independently evolving subpopulations, where the effects of recombination are reduced. Moreover, other phenomena such as selective sweeps and range expansions can generate fitness differentials in sexual populations, potentially contributing to adaptive momentum. While the mechanisms may be more complex in sexual lineages, adaptive momentum remains a plausible explanation for the punctuated evolution observed in the fossil record. Thus, adaptive momentum provides a straightforward and plausible explanation for punctuated evolution and a well-grounded response to assertions that intermittent periods of rapid adaptation must be caused by ill-defined non-Darwinian mechanisms.

## Discussion

We have described and presented evidence for adaptive momentum, a fundamental evolutionary dynamic that has the potential to greatly accelerate the rate of adaptation on multi-peaked adaptive landscapes. We also identified the “balanced” regime, a mode of evolution that exists between the more familiar and well-characterized “sequential fixation” and “clonal interference” regimes. We provide an analytical definition for the balanced regime and identify relevant regions of parameter space, and we demonstrate that this regime provides conditions that allow adaptive momentum to accelerate adaptation.

Prior work has considered how shifts in selection strength over time may affect the mode and tempo of evolution. Indeed, Sewall Wright was the first to discuss macroevolution in terms of fitness landscapes, arguing that “The problem of evolution […] is that of a mechanism by which the species may continually find its way from lower to higher peaks …”^37^. Wright’s Shifting Balance Theory (SBT) was built on the idea that smaller populations experience increased random drift and thus have an enhanced ability to cross fitness valleys. SBT was framed as a three-step process: First, a subpopulation becomes geographically isolated. Then, the subpopulation undergoes drift, leading to the discovery of new adaptive peaks that were otherwise inaccessible. Finally, the subpopulation reintegrates with the main population, allowing the newly discovered beneficial mutations to spread through the entire population. Since its inception, SBT has had its share of supporters and detractors. Among the supporters were Gould and Eldredge, who proposed shifting balance as a mechanism to explain punctuated equilibrium^2^, among other potential mechanisms^38^. However, SBT has often been dismissed, particularly due to problems related to the feasibility of reintegration in the third phase^39,40^. A related theory that sought to understand how increased drift may give rise to the emergence of new species is Mayr’s “founder effect” theory^41^, which posits that speciation typically occurs in small, isolated populations. This theory has also been criticized, because there is scant evidence that new species have experienced such population bottleneck events^42^. More recently, it has been shown that reduced selection at the leading edge of a range expansion can result in the accumulation of deleterious mutations, which can facilitate valley crossings^43^. This reasoning has led to a proposal to extend SBT to include range expansions^44^. Indeed, we consider range expansion to be a mechanism for generating adaptive momentum, because it enhances drift precisely because it weakens selection against deleterious mutations. Thus, we propose an even more comprehensive extension of SBT to include any scenario that produces adaptive momentum, potentially leading to cascades of dynamically coupled adaptations. With this extension, SBT becomes a more powerful potential explanation for variation in rates of adaptation based on shifts in the balance between drift and selection.

We have provided a new mechanism for valley crossings that we call adaptive momentum, and we have shown that its effects on rates of adaptation can be striking in some (but not all) conditions. Many facets of adaptive momentum need to be further explored, such as the effects of different fitness landscapes, the potential interplay among multiple evolving traits, the role of sexual recombination, and the influences of other evolutionary parameters. The effect of the self-imposed constraints of your mathematical fitness landscapes, as well as the consequences of loosening them in future work, are discussed in the Supplementary Discussion section 3. This future work may discover important nuances, complications, and limitations of adaptive momentum. However, we anticipate that the essential dynamics of adaptive momentum will not change.

Simpson identified the greatest hurdle to rapid speciation as the tendency for large populations to experience extreme purifying selection, which he called the “curse of the large population”^1^(p. 95). He argued that the resulting mutation-selection balance in such populations was “unfavorable” for rapid and sustained progressive evolution. However, while Simpson correctly identified that populations at equilibrium are unlikely to discover new fitness peaks, he overlooked the cascading effects of disequilibrium. As we have shown, a population at disequilibrium is poised to exploit adaptive momentum and undergo rapid adaptive evolution. This scenario leaves open the issue of how the state of disequilibrium is initially established, but it can undoubtedly be triggered by various perturbations including environmental instability, population bottlenecks, and the like. Thus, adaptive momentum lifts Simpson’s curse and shows that macroevolution can result from “repeated rounds of microevolution”^45^ after all.

## Online contents

Any methods, additional references, Nature Portfolio reporting summaries, source data, extended data, supplementary information, acknowledgments, peer review information; details of author contributions and competing interests; and statements of data and code availability are available at (http).

## Methods

### Fitness landscape and evolutionary modeling

To isolate the effects of adaptive momentum from other potentially confounding factors, we use a simple tunable fitness landscape that requires multiple consecutive mutations with deleterious effects to cross from one fitness peak to another. (Multiple mutations in a single reproduction event were disallowed in our simulations, see also Supplementary Discussion in Supplementary Information). Individuals are represented as a genome with a single locus holding an integer allele *k* (initially *k* = 0). We model a population of *N* individuals using a Wright-Fisher process. When offspring are produced, the parent’s allele mutates with a probability *µ*, and mutations result in new allele values of *k* +1 or *k* −1 with equal probability. The fitness function is periodic and without diminishing returns (see Figure 2a). The following equations determine how the allele *k* is converted to fitness *w*.

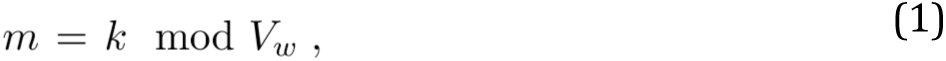

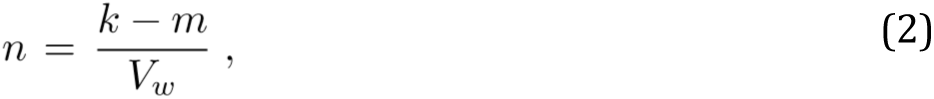

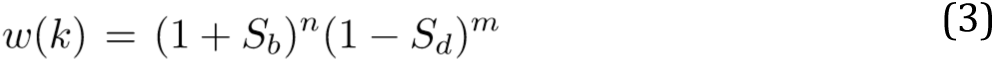

Here, *V_w_* is the width of each valley, *S_b_* is the fitness benefit of each new peak relative to the previous peak, and *S_d_* is the fitness cost of each deleterious mutation in a valley.

The value *m* represents how far a genome has moved across the current valley, and *n* represents the number of peaks already discovered, assuming that the ancestral *k* was initialized to zero. The ascending function *w*(*k*) repeats infinitely and ensures that every valley provides the same challenge and every peak provides the same benefit.

### Parameters for the “three-peaks” experiment

For the “three-peaks” experiment (results in Fig. 2b) we used a version of the landscape described by Eq. (3) that has only three consecutive peaks with *V_w_* = 6, *S_b_* = 0.5, and *S_d_* = 0.03. The population size is *N* = 2,400 and the mutation rate is *µ* = 0.0025 per individual per generation, which produces an average of 6 new mutations per generation in each simulated population. For this experiment, we conducted runs with two different initial conditions. The “fixed start” condition began with all individuals on peak 2. The “mixed start” condition started with 5 individuals on peak 2 and the other 2,395 individuals on peak 1. Each population occupies a 1-dimensional strip of locations with a periodic boundary (i.e., a ring where each individual has two neighbors). Reproduction follows a Wright-Fisher process in which each neighborhood of three individuals (with position indices *n*−1, *n*, and *n*+1) from the current generation (*t*) compete to determine which will produce the individual at the *n*th position in the next generation (*t* + 1). We ran 1000 replicates of each condition for 20,000 generations, and recorded the highest peak that is occupied at the end of each run.

In Extended Data Fig. 1 we show the results obtained using all combinations of ten population sizes (*N* = 400, 600, 800, 1200, 1600, 2400, 3200, 4800, 6400, and 12800) and three costs of deleterious mutations (*S_d_* = 0.024, 0.03, and 0.036). The combination shown at the lower left (*N* = 12800 and *S_d_* = 0.024) maximized the discovery potential. In this condition, the large population size provides a high flux of new mutations, while the shallow valley depth reduces selection against the deleterious mutations needed to discover peak 3. As we move up and to the right, the smaller population sizes and deeper valleys result in fewer discoveries of peak 3, until the fraction of replicates that discover that peak drops almost to 0. Between the extremes, there are conditions where some, but not all, populations discover peak 3; and in these cases, the fraction of replicates that discover peak 3 is usually greater in the mixed-start treatment (orange) than in the fixed-start treatment (black). We can also see a few replicates in the mixed-start runs where populations evidently became stuck on peak 1; in these rare cases, drift evidently caused the extinction of the 5 individuals that started on peak 2 before they could establish a permanent foothold.

### Parameters for the “infinite-peaks” experiment

In the “infinite-peaks” experiment (results in Fig. 3) we measure four statistics related to evolution as a function of population size. Below, we describe the process used to collect the statistics. Extended Data Fig. 2 provides simple illustrations of the statistics and collection methods. Unless otherwise indicated, we use the same parameter values as in the “three peaks” experiment. However, we use an extended version of the fitness function that repeats indefinitely after peak 3. The function is self-similar, such that the benefit from discovering each peak and the difficulty of crossing each valley are identical.

We measured the stochastic tunneling time *T*_st_ by tracking how long an initially fixed population takes to discover and establish at a new peak. A potential discovery is noted when the fitness *w(k)* for any individual in a population reaches a new peak; however, it is only counted as an actual discovery if that value remains at or above the peak for 100 consecutive generations. If a beneficial mutation does not establish, the trial continues, ensuring that *T*_st_ reflects the times for discoveries that actually establish at the new peak, not just transient discoveries that ultimately fail. We measured the neutral drift time *T*_nd_ using the same approach as used to measure *T*_st_, except we set *S_d_* = 0.0. Thus, *T*_nd_ tests the discovery time in the absence of purifying selection. We measured the time it takes a beneficial mutation to sweep to fixation in a population *T*_fix_ by initializing populations with all but one individual on a lower peak and a single individual on an adjacent higher peak. The individuals in the population are each tagged with a neutral genetic marker indicating their starting peak, and we run the simulation until either all individuals have the peak 2 marker, or none did. If the peak 2 marker fixes, then the time of fixation is recorded. If the peak 1 marker fixes, then the trial is discarded. When collecting the *T*_st_, *T*_nd_, and *T*_fix_ data, we always re-initialized the population before starting another test run. Finally, the observed rate of adaptation, *T*_obs_, measures the average time between discoveries of beneficial mutations that become established. We allowed populations to evolve for many generations and recorded the times of each peak discovery, using the same test for establishment as used for *T*_st_.

Because the population sizes varied from 100 to 153,600 in this experiment, the average time between discoveries was highly variable. Thus, it was not feasible to run tests for a fixed number of generations, and instead the tests ran for a fixed computer wall time and used all the resulting data. For *T*_obs_ and *T*_st_, we ran 56 replicates for 12 hours each; for *T*_nd_ and *T*_fix_, we ran 11 replicates for 12 hours each.

Extended Data Fig. 3 and Extended Data Fig. 4 provide additional results generated using the same methods used to generate Fig. 3a. For each panel, we ran a subset of the population sizes from Fig. 3a. In Extended Data Fig. 3, we varied the fitness cost of deleterious mutations *S_d_* and the mutation rate *µ*; in Extended Data Fig. 4, we varied *µ* and the valley width *V_w_*. Changes in all these parameters alter the discovery rates (*T*_st_, *T*_obs_, and *T*_nd_), but not all of them affect fixation times, which depend only on population size and the fitness benefit *S*_b_. These graphs illustrate how the dynamics of evolution depend on the interplay between discovery times and fixation times, illustrating an inverse relation between *S_d_* and *µ* in Extended Data Fig. 3, and between *V_w_* and *µ* in Extended Data Fig. 4. These plots also show that the balanced regime (*T*_nd_ *< T*_fix_ *< T*_st_) consistently accelerates adaptation across these wide ranges of parameter values.

### Logarithmic binning for distribution plots

Fig. 3 shows the distributions *f(T)* of *T*_st_, and *T*_obs_, where *f(T)=n(T)/N_to_*_t_, and *n(T)* is the number of events of duration T that were recorded, and N_tot_ is the total number of events. Because these distributions are highly variable (e.g., for *T*_obs_ values range from <1000 generations at *N* = 153,600 to >10^6^ generations at *N* = 100), we used variable bin sizes to display the distribution, with bin sizes ΔT=0.1T resulting in a constant number of bins on a logarithmic scale. To arrive at a normalized distribution, the probability density function *f(T)ΔT* is plotted in Fig. 3.

### Parameters for the visualization of adaptive momentum

Fig. 4 uses an almost identical setup as the “infinite-peaks” experiment. The only change is that the periodic boundary was disabled. Here, *µ* = 0.0025, *N* = 4800, *V_w_* = 6, *S_b_* = 0.5, and *S_d_* = 0.03.

### Calculation of the index of dispersion

In evolving systems, the dispersion index *R* is used to understand if rates of change depend on Poisson-random event alone or, alternatively, if non-random processes are involved^16^. We quantified *R* for the distributions of *T*_obs_ and *T*_st_ in Fig 3f. To calculate *R*, we first convert the data from times between discoveries to numbers of discoveries per unit of time. As mentioned previously, the time between events (discoveries of a new peak) varies greatly across the test conditions, so we chose a variable unit time of 10 times the mean discovery time for each condition. Thus, on average, there are 10 discoveries per bin. Once we have the counts of discoveries per unit time, *R* is calculated as

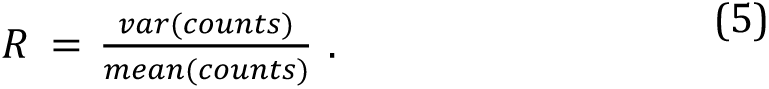

### Evolution of large well-mixed populations

In both the three-peaks and infinite-peaks experiments, the populations evolve in a physically structured environment (specifically, a one-dimensional ring). To determine whether the effects of adaptive momentum also occur in well-mixed populations required the use of a different model because the well-mixed populations have faster fixation times, and thus require larger population sizes to observe effects of adaptive momentum. The larger population sizes could not be supported by the agent-based models described above. Instead, we developed a mathematical simulation based on a stochastic replicator equation^46^. This model tracks the counts of each genotype in the population, then updates the counts at every generation based on each type’s fitness relative to the current average fitness. In our stochastic differential equation we used the counts as the mean parameter in a Poisson distribution and then used a value randomly drawn from that distribution as the actual count, thereby introducing appropriate stochasticity. We then applied mutations, causing shifts in some genotypes to adjacent genotypes in the genomic space. We normalized and discretized the counts every generation to maintain both population size and resolution. We validated our model by comparing its output at lower population sizes to that of the agent-based model, ensuring consistency and accuracy in our results. As shown in Extended Data Fig. 5, adaptive momentum is also seen in well-mixed populations, but the effect of adaptive momentum manifests differently, as discussed in Supplementary Information (Supplementary Discussion, section 2).

## Acknowledgements

We thank Jonathan Lansey, Matt Rupp, and Thomas LaBar for contributions to an earlier version of this study that referred to the “Free-for-All Effect”. This work was supported in part by Michigan State University through computational resources provided by the Institute for Cyber-Enabled Research., and by the National Science Foundation under Cooperative Agreement No. DBI-0939454.

## Extended Data Figures

**Extended Data Fig. 1.**
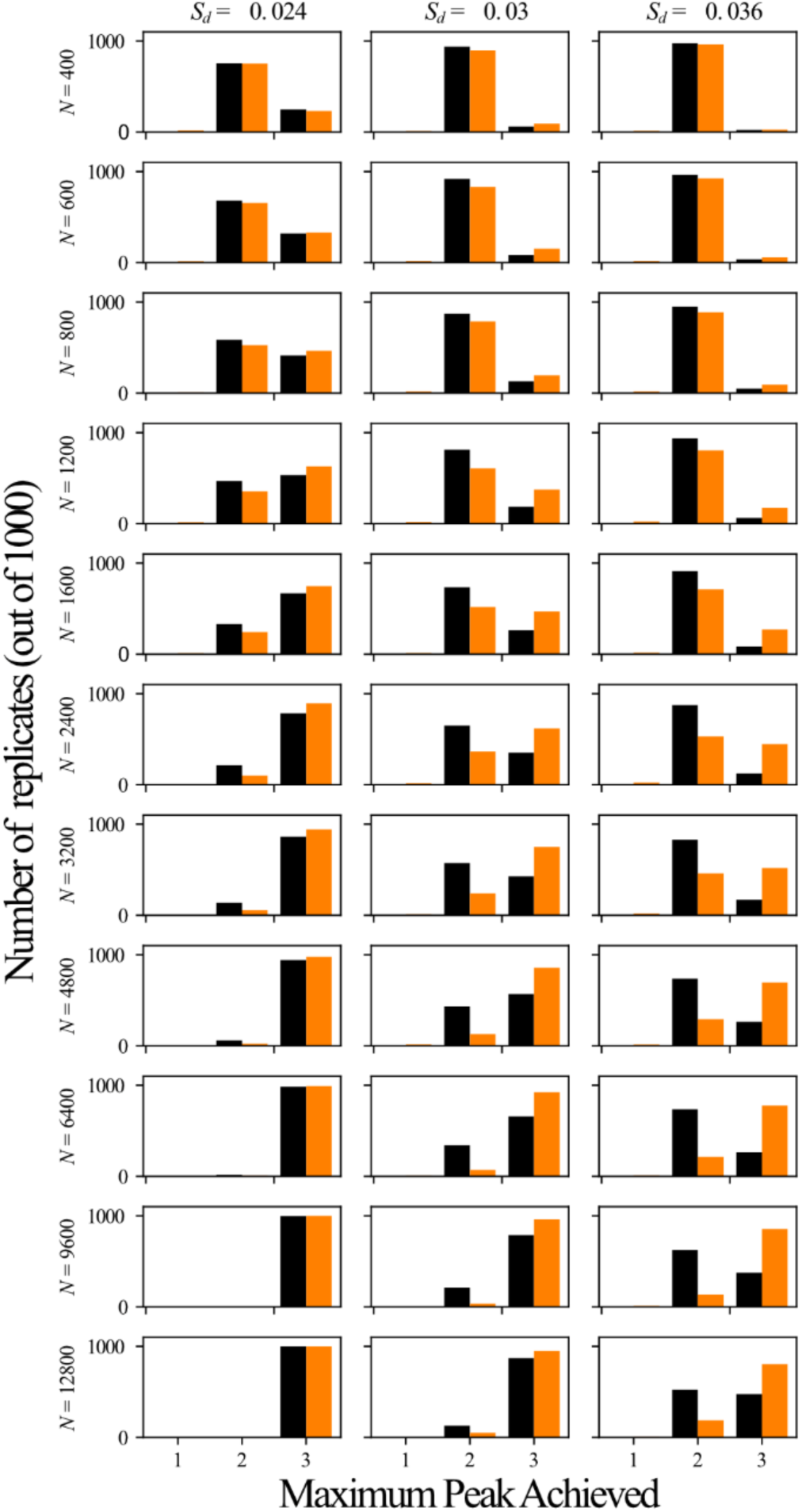
Histograms of highest peak achieved in the “three-peaks” experiment using different valley depths and population sizes. Black bars show the final counts for the fixed-start condition. Orange bars show the final counts for the mixed-start condition. Each row of panels used a different population size, increasing from *N* = 400 at the top to *N* = 12800 at the bottom. Each column used a different valley depth, increasing from *Sd* = 0.024 at the left to *Sd* = 0.036 at the right.

**Extended Data Fig. 2.**
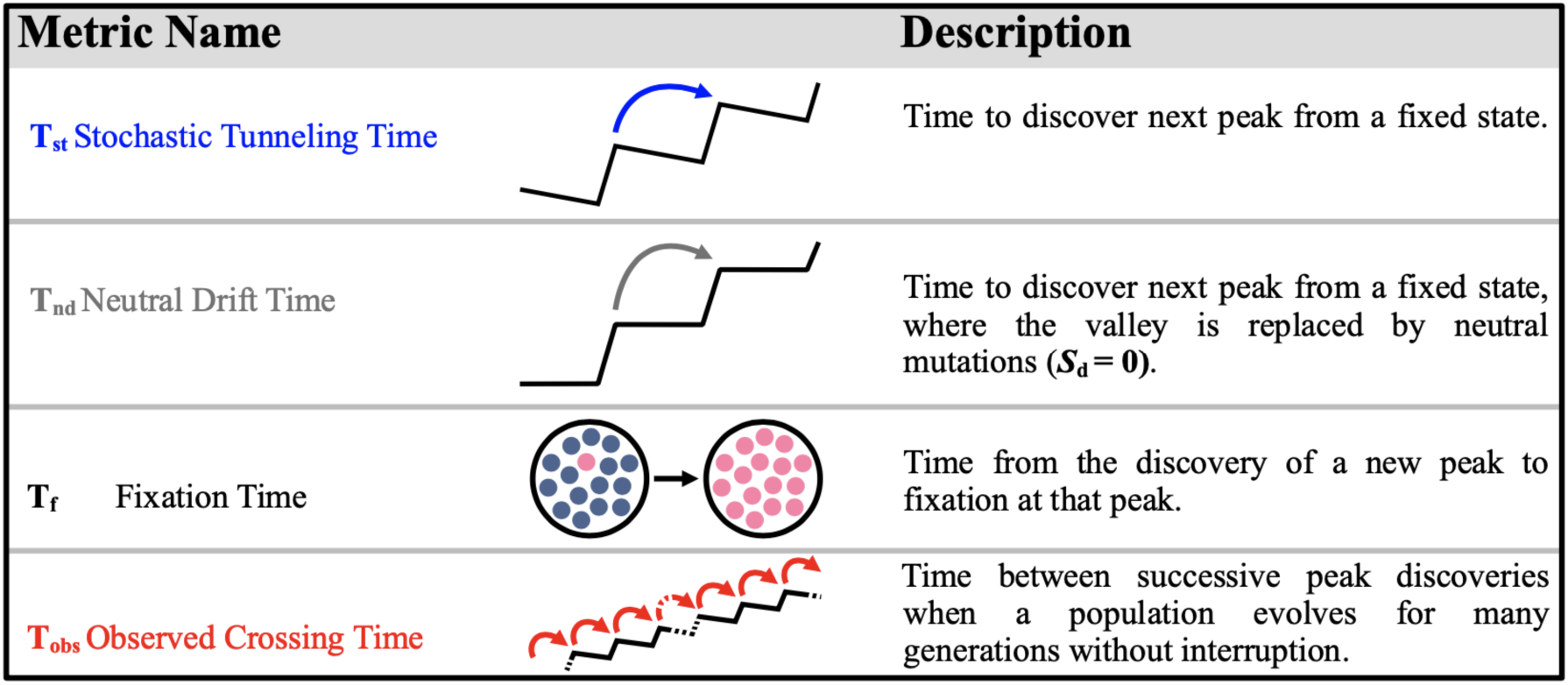
Schematic illustration of Tobs, Tst, Tnd, and Tfix.

**Extended Data Fig. 3.**
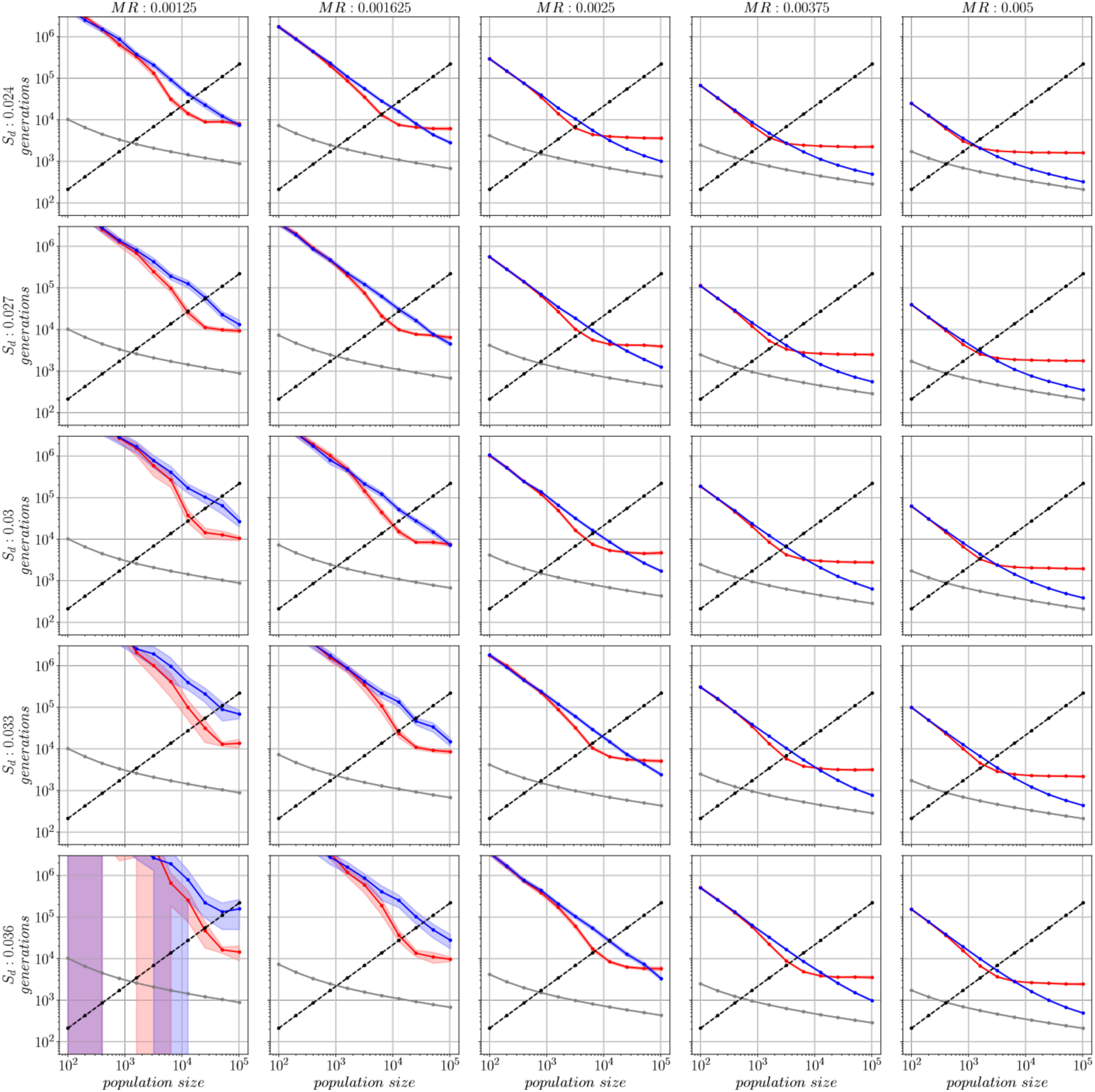
Average valley-crossing times and fixation time as a function of mutation rate *µ* and valley depth *Sd* in the “infinite peaks” experiment. Each panel shows the average stochastic tunneling time *T*st (blue), neutral drift time *T*nd (gray), fixation time *T*fix (black), and observed crossing time *T*obs (red) as a function of population size for the fitness function described in Eq. 3 and the indicated deleterious mutation costs and mutation rates. Each column used a different mutation rate, increasing from *µ* = 0.00125 (left) to *µ* = 0.005 (right). Each row used a different deleterious mutation cost, increasing from *Sd* = 0.024 (top) to *Sd* = 0.036 (bottom). The mutation rate *µ* = 0.0025 and deleterious mutation cost *Sd* = 0.03 in the center panel reproduce the data in Fig. 3a (main text). More costly values of *Sd* and lower mutation rates make the discovery of beneficial mutations less likely, leading to a tradeoff between these parameters that is manifest by the similarities along the diagonals from the upper-left to lower-right panels.

**Extended Data Fig. 4.**
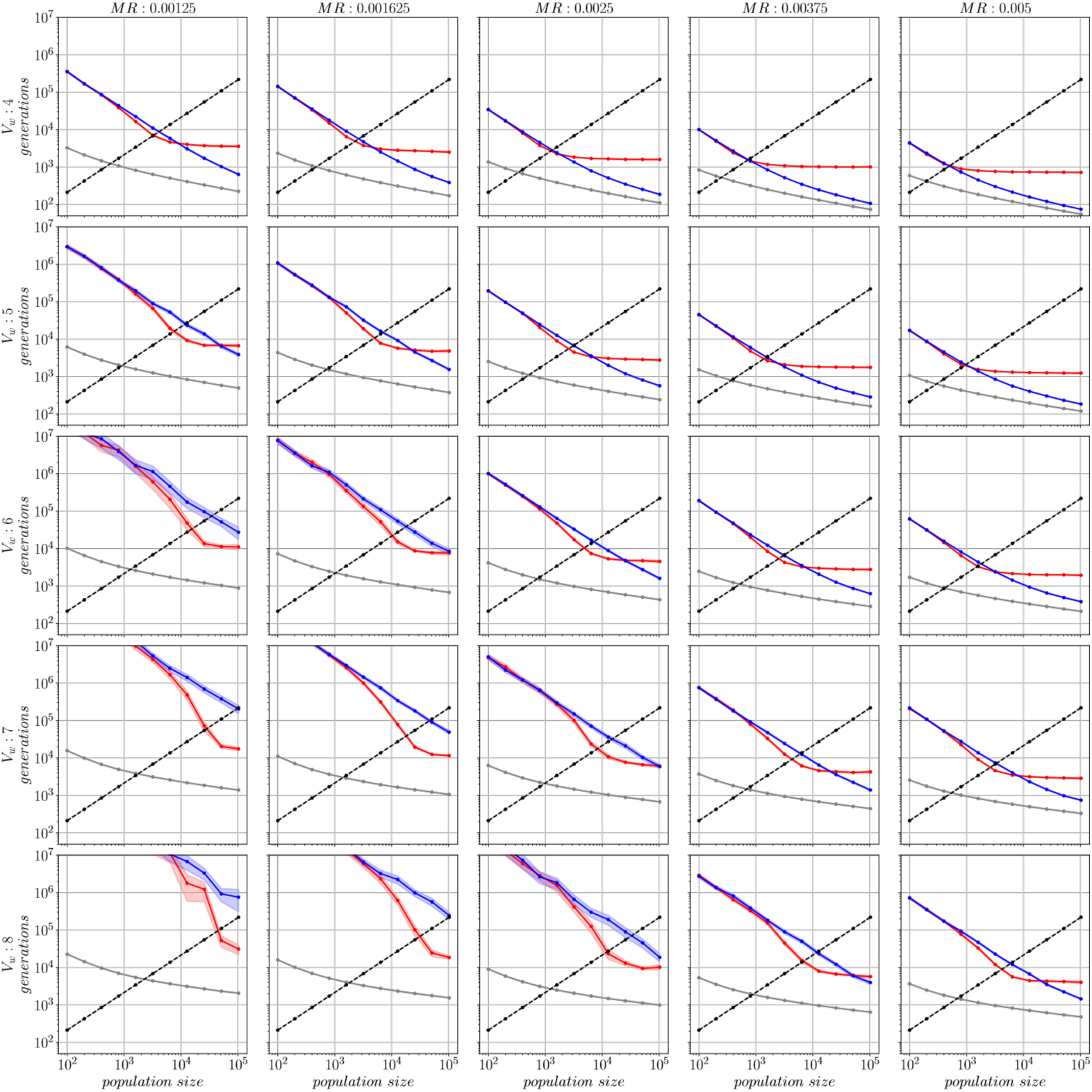
Average valley-crossing times and fixation times as a function of mutation rate *µ* and valley width *Vw* in the “infinite peaks” experiment. Each panel shows the average stochastic tunneling time *T*st (blue), neutral drift time *T*nd (gray), fixation time *T*fix (black), and observed crossing time *T*obs (red) as a function of population size for the fitness function described in Eq. 3 and the indicated valley widths and mutation rates. Each column used a different mutation rate, increasing from *µ* = 0.00125 (left) to *µ* = 0.005 (right). Each row used a different valley width increasing from *Vw* = 4 Top) to *Vw* = 8 (bottom). The mutation rate *µ* = 0.0025 and valley width *Vw* = 6 in the center panel reproduce the data in Fig. 3a (main text).

**Extended Data Fig. 5.**
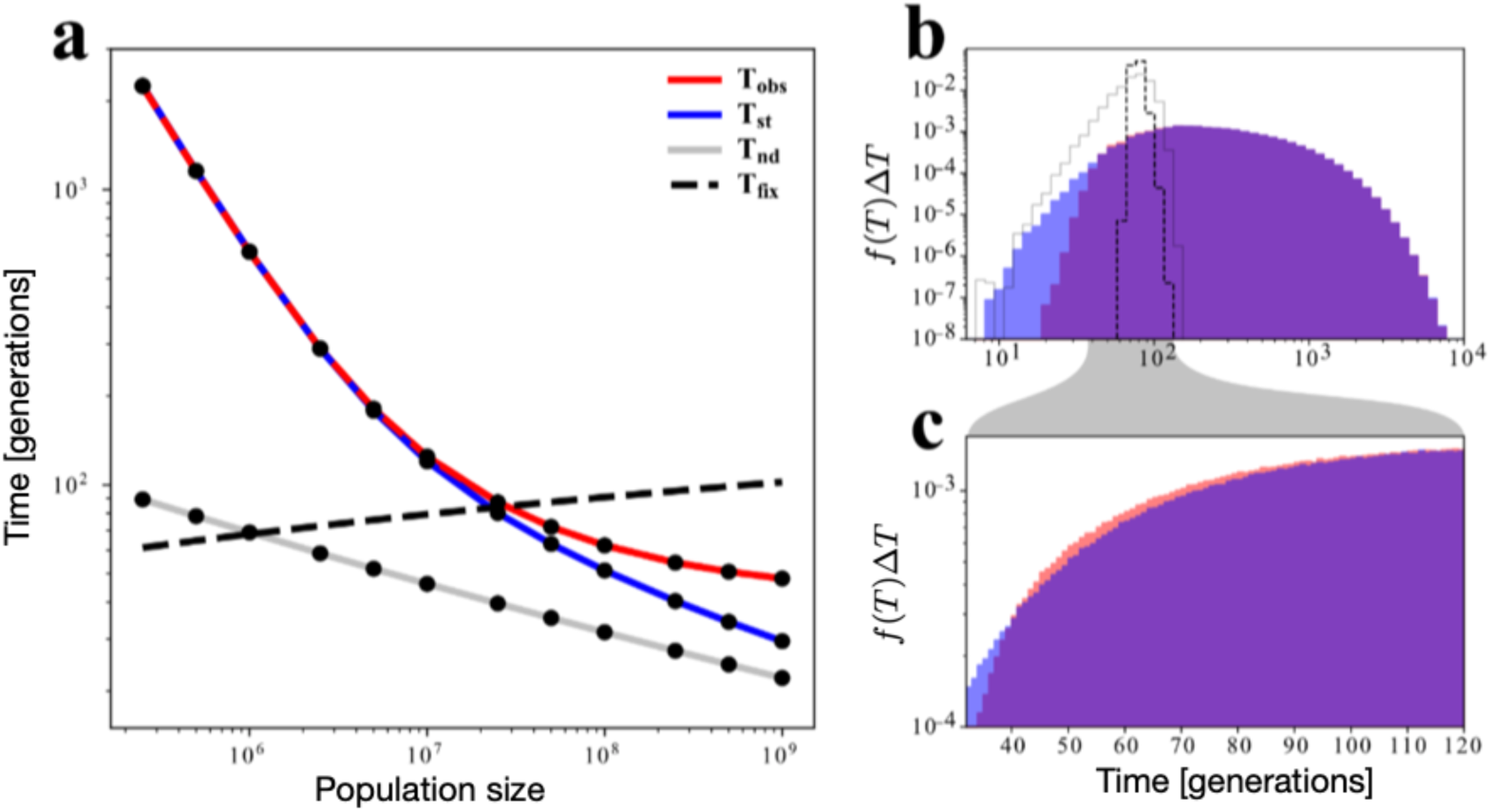
Average valley-crossing and fixation times and distributions in the “infinite-peaks” experiment for well-mixed populations. **a**, Average stochastic tunneling time *T*st (blue), neutral drift time *T*nd (gray), fixation time *T*fix (black), and observed crossing time *T*obs (red) as a function of population size for the fitness function described in Eq. (3) and shown in Fig. 2a. **b,** Distributions *f(T)* of *T*st (blue), *T*obs (red), *T*nd (black-dashed outline), and *T*fix (gray-solid outline) for *N* = 10^9^. Generations are binned on a logarithmic scale. **c**, Detailed view of the *T*obs and *T*st distributions shown in **b**, highlighting the distributions between *T*=30 and *T*=120. In the detailed view, generations are binned on a linear scale.

## Supplementary Information

### Supplementary Discussion

#### 1. Adaptive momentum is modulated by population structure

Adaptive momentum can be understood from fundamental principles, without reference to the population structure. However, population structure can significantly affect evolution^47^, and thus adaptive momentum, as well. In particular, population structure affects fixation times^48^ and population structure influences which individuals in a population compete against each other and thus which individuals experience adaptive momentum, when they experience it, and to what extent.

As discussed in the main text, fixation time is critical to determining if adaptive momentum is likely to accelerate evolutionary outcomes. In highly connected populations, a beneficial genotype will spread more quickly than in populations with sparse connectivity, resulting in shorter fixation times. As a result, in well-mixed populations where every individual competes directly with every other individual (i.e., no spatial structure), momentum windows are likely to be very short and thus inhibit adaptive cascades. Conversely, spatially structured populations will tend to have longer fixation times (inversely related to the degree of connectivity), which lengthen momentum windows, making cascades more likely.

In addition to affecting fixation times, population structure also affects the dynamics of selection. In a spatially structured population, reproductive competition occurs within neighborhoods local to each individual. After the initiation of disequilibrium (e.g., an initial discovery and establishment of a beneficial mutation), the subset of the population with the beneficial alleles will grow outward from the point of initiation. This growth results in a leading-edge interface comprised of individuals on one side with relatively high fitness competing against lower-fitness individuals on the other side. The number of individuals along (or near) the expanding leading edge, and thus the number of individuals that experience adaptive momentum, depend on the shape of that leading edge, which in turn can depend on geographic features of the environment and other elements that affect population connectivity.

Beyond simply having elevated fitness and thus an elevated capacity to buffer mutations, Individuals at the leading edge will be the offspring of parents who were previously on the leading edge. Thus, in spatially structured populations expanding subpopulations generate propagating lineages comprised of individuals who are the result of consecutive generations of reduced selection. The leading-edge subpopulations are protected from the harshest effects of purifying selection, and thus they can become hotbeds of innovation, able to explore genetic states that may be inaccessible to individuals not on the leading edge who must contend with purifying selection^49^. However, individuals within protected lineages are not immune from selection: if a lineage accumulates mutations that are too costly, they may be overtaken by less encumbered lineages.

Finally, in spatially structured populations, there tends to be a relatively constant ratio of advantaged and disadvantaged individuals within local neighborhoods along the leading edge. As a consequence, there tends also to be a relatively constant elevation in the potential for leading-edge lineages to cross valleys and discover new fitness peaks during a momentum window.

The dynamics of selection in well-mixed populations are quite different. In well-mixed populations, every individual competes directly with every other individual, and so there are no leading edges that can localize the effects of adaptive momentum. In other words, there is no leading-edge subpopulation with a maintained reduction in selection pressure as in spatial populations. Rather, in well-mixed populations the advantage from adaptive momentum is neither localized to a subset of advantaged individuals nor constant over time. Instead, the entire subpopulation of advantaged individuals experiences reduced selection pressure based on the magnitude of their advantage relative to the mean fitness of the population. As the population equilibrates, the population’s average fitness approaches the fitness level of the most advantaged individuals (i.e., those without any deleterious mutations), and the exploratory advantage provided by adaptive momentum decays. This equilibration process results in a variable selective advantage over a momentum window, one that starts strong but then decays over time.

In addition to a variable strength of selection, in well-mixed populations advantaged subpopulations experience variability in mutation supply during momentum windows. Because all individuals with the advantage experience reduced selection from adaptive momentum, and because this subpopulation grows during a momentum window, the mutation supply available to this subpopulation increases as the number of individuals with the advantage increases. The mitigating effect of a growing advantaged clade also exists in spatial populations, but here the slower fixation rates and limited effective area of the leading edge blunt the mitigating effect (particularly in the one-dimensional populations used in the main paper results).

In summary, the effect of adaptive momentum in well-mixed populations is variable over time. Its effect on individuals should be strong soon after a beneficial mutation occurs, but this effect is mitigated by the limited number of advantaged individuals, constraining mutation supply and beneficial mutation discovery. As the advantaged individuals proliferate, the increased mutation supply increases the potential for valley crossings and discovery of new peaks. However, as the advantaged lineage becomes common, the population’s average fitness also increases, resulting in stronger purifying selection and a diminished capacity for exploration.

The data in Figure 3 was generated using a spatial one-dimensional “ring” population structure. That spatial structure maximizes the fixation time of beneficial mutations, and thus it also maximizes the potential for adaptive cascades. In the next section, we describe results from an experiment using a well-mixed population structure (with Wright-Fisher selection). That experiment shows that, while adaptive momentum is much harder to detect and quantify, it is still present.

#### 2. Adaptive momentum in unstructured populations

Extended Data Figure 5 presents data from an experiment using well-mixed populations. The experiment uses the same fitness function and parameters as Figure 3. However, we analyzed larger population sizes due to the short fixation times typical of well-mixed populations, where the intersections of *T*_fix_ with *T*_nd_ and *T*_st_ occur at larger population sizes. In Extended Data Figure 5a, we can see patterns indicative of both sequential fixation (SF) and clonal interference (CI) regimes. However, unlike in Figure 3, there is no discernible offset between *T*_st_ and *T*_obs_ in the balanced regime; that offset would illustrate accelerated evolution due to adaptive momentum. Nonetheless, a closer examination of the distributions of *T*_st_ and *T*_obs_ at a population size of 1,000,000 reveals a robust increase in the frequency of observed discovery times in the *T*_obs_ distribution between 40 and 120 generations in comparison to the *T*_st_ distribution (Extended Data Figure 5b-c). This increase does not translate into a visible offset for the averages shown in Extended Data Figure 5a, however, owing in part to the very small magnitude of the difference between the distributions, but also because of a “refractory effect”, a phenomenon we now describe.

Following the appearance of a new beneficial mutation, it is necessary for a critical mass of the new variant to accumulate before subsequent beneficial discoveries become statistically likely. While the new variant enjoys a high selective advantage at the onset of a sweep, the new variant is still rare, so, the mutation supply is insufficient for effective exploration. As a consequence, the distribution of *T*_obs_ is missing short transition times that are part of the *T*_st_ distribution in Extended Data Figure 5b. The low mutation supply condition during the early stages of a selective sweep should delay the discovery of new peaks under all conditions, regardless of population structure. However, because well-mixed populations tend to have shorter fixation times and any discoveries of new peaks produced by adaptive momentum must occur within that short window, well-mixed conditions also tend to magnify the effect of the refractory period.

The tail with short transition times in the *T*_st_ distribution (that is missing in the *T*_obs_ of well-mixed populations) exists because our procedure for measuring stochastic tunneling times *T*_st_ starts with the entire population on one fitness peak, which is immediately available for tunneling to the next peak and upon discovery and establishment, the population is reset before another measurement is made. By contrast, in the *T*_obs_ data, the populations are not reset after each beneficial mutation discovery. Thus, the *T*_obs_ distribution times are offset by the time necessary for each advantaged subpopulation to grow to a large enough size that the influx of mutations is sufficient for valley crossing to the next peak. In Extended Data Figure 5, the refractory period takes up a large fraction of the momentum window (Extended Data Figure 5b and c). Conversely, spatially structured populations exhibit longer fixation times, and the impact of the refractory period is substantially diminished.

#### 3. Effect of constraints on the fitness landscapes and future directions

The fitness landscape (as well as the evolutionary dynamics on it) that we used to obtain our results is necessarily abstract. In this section we discuss how releasing some of the constraints might affect adaptive momentum.

As described in Methods, each newly produced offspring can gain only a single mutation. We imposed this constraint to prevent rare valley-crossing events produced by multiple mutations, which might complicate interpretation. Allowing multiple mutations should facilitate valley crossing and thereby shorten discovery times. Implementing multiple mutations would slightly change the relations between the drift, tunneling, fixation and observed discovery times. It might shift the conditions (e.g., population size) where adaptive momentum has the greatest effect and the boundaries of the balanced region, but it would not fundamentally change the dynamics of adaptive momentum. We used a regular (i.e., repeating) fitness function that allowed us to gather meaningful statistics about valley crossings from a single evolutionary run. If we had used an irregular fitness function instead, it would be difficult to distinguish valley crossings that are short due to adaptive momentum from those that are short because a particular valley happened to be easy to cross.

Another feature of our landscape is the absence of diminishing-returns epistasis, which is a feature of many fitness landscapes^50–54^. This form of epistasis causes a slowing of adaptation over time because each successive benefit tends to be harder to discover and less beneficial than the last. In an extreme case, a landscape with diminishing returns may exhibit adaptive momentum only after an initial phase of clonal interference during which beneficial mutations are easily discovered. However, this period might eventually end as the population becomes well-adapted, beneficial mutations become rare, and sequential fixation sets in. Despite this potentially reduced efficacy over time, the fundamental dynamics of adaptive momentum would not be altered.

In this work, we have examined adaptation based on only a single organismal trait described by the fitness function. In reality, organisms have a multitude of traits that interact to produce their overall fitness. Future research should explore multi-trait fitness functions^55^, where multiple adaptive directions exist at every point in time. Rather than modeling a one-dimensional fitness landscape with peaks and valley, we could model a more complex landscape involving peaks and valleys in independent dimensions and combine those values into an overall fitness with or without epistasis between traits. We hypothesize that adaptive momentum based on progress in one trait will reduce selection pressure across all traits, perhaps facilitating concurrent improvement in multiple traits and potentially allowing the crossing of even wider and deeper fitness valleys.

Another consequence of multi-trait fitness functions is the possibility of gene interactions: the condition in which the allele of one gene alters the fitness effects of mutations on other genes (epistasis, see e.g.,^56^). While the system we use does have sign epistasis between mutations, it does not include the possibility of more complex interactions between traits that may occur on higher-dimension fitness functions. While a system with gene interactions will still experience adaptive momentum, it will be difficult to distinguish short crossing times resulting from adaptive momentum from short crossing times resulting from gene interactions.

Yet another future direction would explore the effects of sexual recombination on the dynamics of valley crossings and the role of adaptive momentum. On one hand, recombination can more effectively purge deleterious mutations, making valley crossing less likely^57,58^. On the other hand, recombination can accelerate the production of combinations of deleterious mutations, potentially enhancing adaptive momentum as well as stochastic tunneling^59^. Furthermore, recombination can alleviate clonal interference^60,61^, potentially extending the effects of adaptive momentum to larger populations. In other words, the rate of adaptation might not be as impacted when *T*_fix_ > *T*_st_ because recombination would resolve some instances of clonal interference. Future research is warranted on all these effects. The addition of sexual selection may have drastic effects when integrated into adaptive momentum theory as any shift in preference may result in adaptive momentum until a newly preferred trait becomes common^55^.

